# Sensitive and reproducible determination of clinical HDL proteotypes

**DOI:** 10.1101/2020.07.09.191312

**Authors:** Sandra Goetze, Kathrin Frey, Lucia Rohrer, Silvija Radosavljevic, Jan Krützfeldt, Ulf Landmesser, Marco Bueter, Patrick G. A. Pedrioli, Arnold von Eckardstein, Bernd Wollscheid

## Abstract

**Background:** High-density lipoprotein (HDL) is a heterogenous mixture of blood-circulating multimolecular particles containing many different proteins, lipids, and RNAs. Recent advancements in mass spectrometry-based proteotype analysis strategies enable the sensitive and reproducible quantification of proteins across large patient cohorts.

**Methods:** HDL particles were isolated from plasma of more than 300 healthy individuals or patients with a multiplicity of physiological HDL states. From these, peptides were extracted and HDL proteome spectral libraries were generated. This is a prerequisite for using data-independent acquisition (DIA) strategies to analyze HDL particles from clinical cohorts using mass spectrometry.

**Results:** The resulting HDL proteome spectral libraries consist of 296 protein groups and 341 peptidoforms of potential biological significance identified with high confidence. We used the HDL proteome libraries to evaluate HDL proteotype differences in between healthy individuals and patients suffering from diabetes mellitus type 2 (T2DM) and/or coronary heart disease (CHD). Bioinformatic interrogation of the data revealed significant quantitative differences in the HDL proteotypes including a significant depletion of phosphatidylinositol-glycan-specific phospholipase D (PHLD) from disease-derived HDL particles.

**Conclusion:** The DIA-based HDL proteotyping strategy enabled sensitive and reproducible digitization of HDL proteotypes derived from patient cohorts and provides new insights into the composition of HDL particles as a rational basis to decode structure-function-disease relationships of HDL.

**List of human genes and protein names discussed in the paper:**

- APOA1 (Apolipoprotein A-I)
- APOA2 (Apolipoprotein A-II)
- APOE (Apolipoprotein E)
- APOC3 (Apolipoprotein C3)
- CLUS (Clusterin)
- PHLD (Phosphatidylinositol-glycan-specific phospholipase D)
- PON1 (Serum paraoxonase/arylesterase 1)
- PON3 (Serum paraoxonase/lactonase 3)
- PSPB (Pulmonary surfactant-associated protein B)
- RAB1B (Ras-related protein Rab-1B)
- RAB6A (Ras-related protein Rab-6A)
- RB11A/B (Ras-related protein Rab-11A/B)
- RP1BL (Ras-related protein Rap-1b-like protein)
- RAB10 (Ras-related protein Rab-10)
- SAA1 (Serum amyloid A-1 protein)
- SAA2 (Serum amyloid A-2 protein)
- SAA4 (Serum amyloid A-4 protein)
- SCRB1 (Scavenger receptor class B member 1)

## Introduction

High-density lipoprotein (HDL) is the smallest and densest of all lipoprotein particles in the human body. It contains the highest proportion of proteins relative to lipids of characterized lipoprotein particles. The most abundant apolipoproteins in the particle are APOA-I and APOA-II, which account for about 65% and 15%, respectively, of the protein mass in HDL particles (1). HDLs exert a multiplicity of functions potentially beneficial for human health. It is mostly known for mediating reverse cholesterol transport from peripheral tissues and arterial walls back to the liver, either specifically via the interaction with scavenger receptor class B member 1 (SCRB1) and ATP binding cassette (ABC) transporters or nonspecifically by aqueous diffusion (1). Changes in cellular cholesterol homeostasis have been associated with anti-inflammatory and anti-oxidative effects of HDL and with increases in endothelial nitric oxide synthase activity that induce production of the anti-atherogenic molecule nitric oxide (NO) (2). In addition to these global effects, HDL particles can induce signaling processes in cells through specific ligand-receptor interactions (2).

Low plasma levels of HDL cholesterol (HDL-C) are associated with higher risk of several diseases including coronary heart disease (CHD) and diabetes mellitus type 2 (T2DM) (3). However, the cholesterol content of HDL particles does not seem to be causative of disease development. Rather, cholesterol appears to be a non-functional constituent of HDLs that clinically serves as an approximation for HDL particle number and size but does not provide insights into particle heterogeneity or functionality (4). In the past decade, it has become clear that the HDL proteome is complex ((5–25) and (http://homepages.uc.edu/~davidswm/HDLproteome.html)) and not uniformly distributed among HDL subclasses (26). These differentially distributed proteoforms can affect lipid metabolism as well as various biological processes such as complement activation or the acute-phase immune response (27). Our understanding of structure-function relationships of HDL is limited in large part due to our incomplete knowledge of identities and quantities of HDL constituents.

Here, we set out to characterize the HDL proteotype using next-generation mass spectrometry-based analysis strategies. In contrast to data-dependent acquisition (DDA) strategies, data-independent acquisition (DIA) strategies for mass spectrum (MS) analyses enable the consistent recording of fragment ions from all peptide precursors/proteins present in a biological sample (28). DIA-MS using spectral libraries therefore allows for more reproducible and accurate protein identification and quantification than does DDA-MS. The DIA-MS strategy enables not only the digitization of patient cohorts but also multi-centric data generation and data analysis/interoperability (29). The spectral libraries generated by us include 296 protein groups and 341 peptidoforms (i.e., post-translationally modified peptides) of potential clinical significance, enabling sensitive and reproducible DIA-MS HDL proteotyping across patient cohorts. We used this HDL proteotyping strategy to quantitatively interrogate HDL proteotype differences between healthy subjects and T2DM and CHD patient cohorts. Quantitative HDL proteotype differences included a significant depletion of phosphatidylinositol-glycan-specific phospholipase D (PHLD) from disease-derived HDL particles. The toolbox and strategy developed in combination with the data generated will serve as a rational basis for further elucidation of HDL protein structure-function relationships to support clinical decision making.

## Materials and Methods

### Clinical HDL sample collection

The Institute of Clinical Chemistry of the University Hospital Zurich (USZ) provided plasma pools from healthy individuals as well as subjects with T2DM (defined as fasting glucose >8 mmol/L to make outliers of otherwise euglycemic subjects unlikely), CHD (according to the guidelines of the American Heart Association), the combination of T2DM and CHD, acute inflammation (defined by C-reactive protein >200 mg/L), and renal failure (defined as eGFR <40 mL/min/1.73 m2) were included. In addition, we included pools of dyslipidemic plasmas with either HDL-C <1 mmol, triglycerides >4 mmol/L, or total cholesterol >7.5 mmol/L. Additionally, samples from obese patients, taken prior to and post bariatric surgery, were included. Ethics approvals were obtained from Kantonale Ethikkommission Zurich (PB-BASEC_2015-00159 and PB-BASEC_2016-02287) or Ethikkommission der Charité – Universitätsmedizin Berlin (EA4/123/16). All subjects gave informed consent.

### Clinical sample preparation for HDL proteotyping

HDL particles were isolated by differential ultracentrifugation. Serum density was adjusted to 1.063 with KBr. The top LDL-containing layer was removed, and the density was raised to 1.21 with KBr. The top HDL-containing layer was removed and delipidated using methanol/chloroform extraction. Proteins were reduced, alkylated, and digested using sequencing-grade trypsin (Promega) in a ratio of 1:40 (see Supplemental Materials and Methods for details).

### Spectral library generation for HDL proteotyping

Peptides were separated by reversed-phase chromatography on a high-pressure liquid chromatography column (EASY-Spray RSLC C18, 2 *μ*m, 50 cm x 75 *μ*m) (Thermo Fisher Scientific), which was connected to a nano-flow HPLC interfaced with an autosampler (EASY-nLC 1200, Proxeon). The HPLC was coupled to a QExactive HF (Thermo Fisher Scientific) equipped with an Easy-Spray ion source (Thermo Fisher Scientific). Peptides were separated and mass spectra were acquired as described in Supplemental Materials and Methods. Raw files, processed data, and spectral libraries are available via the public MS data repository Massive with the identifier MSV000084733.

For HDL spectral library generation, DDA raw files were processed with Proteome Discoverer software version 2.2 using a human UniProt database (release 201804) together with iRT peptides (Biognosys) and common contaminants. Two spectral libraries were generated: one including standard peptide modifications and a second with biologically relevant post-translational modifications (PTMs). To identify PTMs, we first performed an open search using MSFragger (30). From this we selected the eight most prevalent modifications (methionine oxidation, tryptophan oxidation, serine, threonine, and tyrosine phosphorylation, lysine mono-and dimethylation, and protein N-terminal acetylation) and performed a closed search in Proteome Discoverer 2.2 (for details see Supplemental Materials and Methods). Both libraries were assembled in Spectronaut v13 (Biognosys) using standard parameters. The identifications were filtered for 1% FDR on peptide and 5% FDR on protein level. The spectral library for DIA analysis contained 428 unique protein groups and 5195 peptide precursors. The spectral library for DIA peptidoform analysis contained 437 unique protein groups and 5556 peptide precursors. For Gene Ontology (GO) analysis (David GO direct) our standard spectral library was applied. The p-value was determined with the Fisher’s exact test and the significantly enriched biological processes (p-value *≤* 0.05) were grouped into nine umbrella terms describing known HDL functions (27).

### DIA-MS HDL proteotyping of clinical cohort

For the quantitative DIA-MS HDL proteotyping analysis, samples from 51 healthy volunteers, 46 patients with T2DM, 25 with CHD, and 19 patients with both T2DM and CHD were recruited at the University Hospital Zurich or the Charité Universitätsmedizin Berlin (for details of the patient cohort see (3)). DIA data processing is described in Supplemental Materials and Methods. For statistical data evaluation, MSstats3 (v3.12.3) was used (31) (see Supplemental Materials and Methods for details). Data was processed in R (version 3.6.1.) and visualized with the ggplot2, gplots, and fmsb packages. For modified peptide selection and feature clustering, the quantitative peptide matrix generated by Spectronaut v13 based on the peptidoform library with additional PTMs was used after removing mass spectrometry contaminants and non-proteotypic peptides. Intensities were log10 converted, and an intensity-filtered quantitative matrix was used (see Supplemental Materials and Methods for details). Using an ExtraTreesClassifier estimator from the scikit-learn python library, the 10 most important features for health versus disease were repeatedly selected (scikit-learn RepeatedStratifiedKFold, 2 repeats and 6 folds) using recursive feature elimination as previously described (32). Features (i.e., proteins, peptides, and modified peptide ratios) selected in more than half of the cross-validation runs were used to generate the clustermaps.

## Results

### DEVELOPMENT OF AN HDL PROTEOME SPECTRAL LIBRARY

The sensitive and reproducible identification and quantification of proteins and their post-translational modifications across clinical sample cohorts is a prerequisite for the functional analysis of biological processes and to link structure with function. Here, we set out to build tools to enable the characterization of HDL on the proteotype level. For discovery-driven generation of HDL proteome spectral libraries, we analyzed pooled patient samples from more than 300 HDL isolates representing a multiplicity of HDL states via 106 DDA-MS experiments. HDL preparations were derived from healthy individuals and patients suffering from inflammation, kidney failure, impaired renal function, low levels of HDL-C, hypertriglyceridemia, hypercholesterolemia, T2DM, and CHD. Additionally, samples from obese patients taken prior to and post bariatric surgery were included in the discovery-driven DDA-MS experiments. The DDA-MS-based strategy established an HDL spectral library encompassing 428 individual protein groups (**Table S1**). The initially identified protein groups were filtered for known contaminants such as, skin-associated proteins (e.g., keratins, potentially introduced by blood withdrawal and sample processing), histones, and APOB and Apo(a) (by current definition part of LDL or Lp(a)). This filtering resulted in a high-confidence HDL proteome spectral library consisting of 296 different protein groups (**Fig. 1**, red circle), containing proteins that were identified at least twice. Proteins detected only once within our DDA-MS measurements were included in the library if literature-based evidence supported inclusion.

**Fig. 1.**
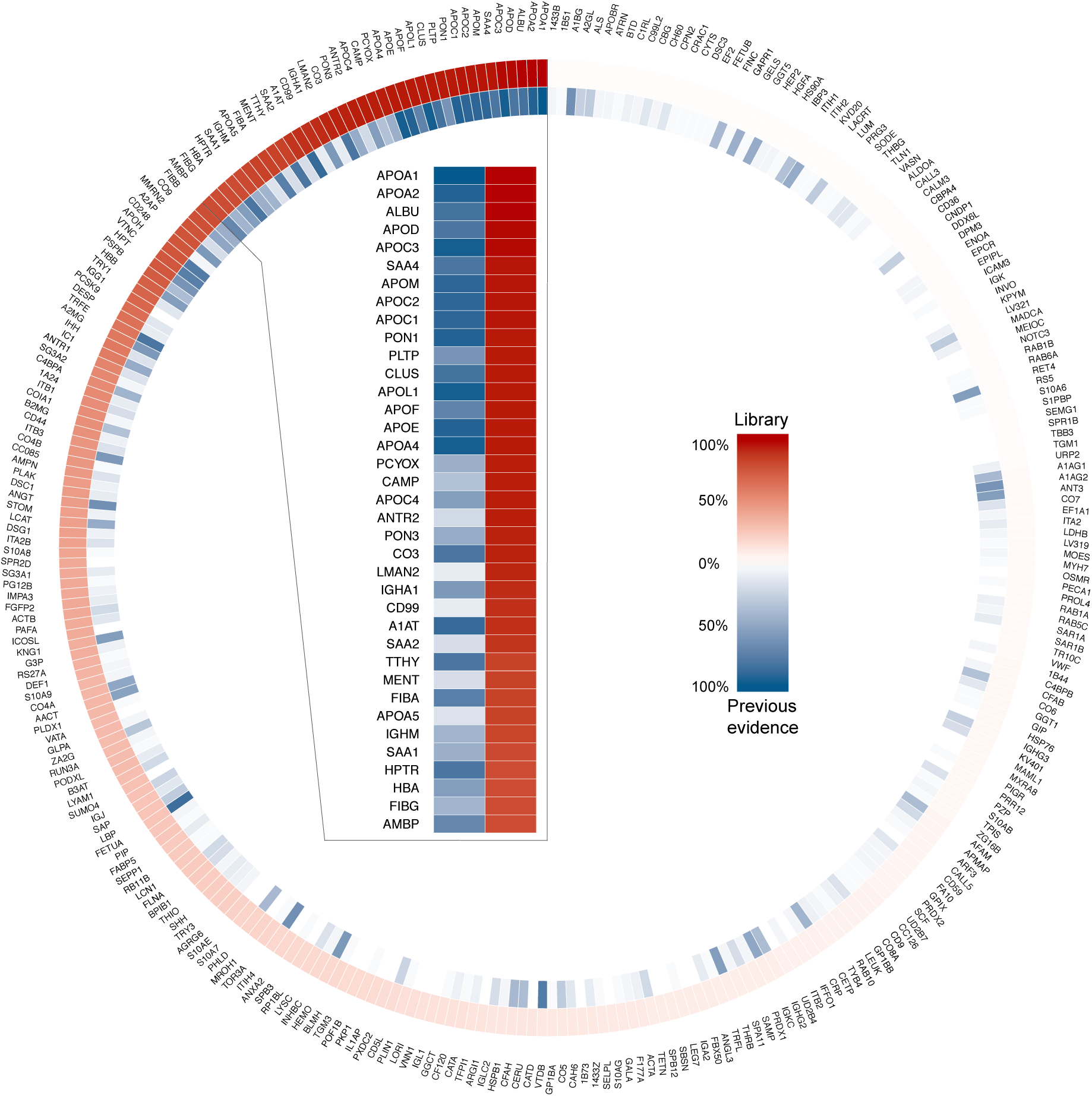
Overview of protein constituents of HDL identified in our analysis and previous studies. The high-confidence HDL proteome contains 296 protein groups. In red are shown proteins detected in our analysis at least twice or once if supported by prior literature. In blue are shown literature-derived HDL proteins. Only proteins overlapping with our HDL proteome spectral library are depicted. Color gradients indicate protein group identification frequency. Proteins identified in more than 80% of our sample pool are highlighted in the insert.

We then compared our HDL proteome spectral library to previously published datasets on HDL proteome constituents including the Davidson HDL watchlist (http://homepages.uc.edu/~davidswm/HDLproteome.html, last updated August 2015) and 20 HDL proteome analysis studies published between 2015 and February 2020 that included protein annotation data (5–23,25). In most of these studies HDL was isolated using an ultracentrifugation protocol similar to our HDL isolation strategy. Recent data sets derived from size-exclusion chromatography (10,11,18,19,21) or affinity purification were included in the comparison (6,7,25). In total, this led to a list of 223 literature-derived protein groups that were identified in three or more independent studies (**Fig. 1**, blue circle, **Table 1, Table S2**).

**Table 1.** Integrated summary of HDL protein identifications across different resources. The table provides an overview of the frequency of identified protein groups in HDL proteome spectral library (red) and relates these protein groups to formerly identified HDL proteome constituents described in other studies (blue).

| Number of identifications |  | Protein Groups |
| --- | --- | --- |
| HDL library | Independent studies |  |
| $\geq 1$ | $\geq 3$ | **150** |
| $\geq 2$ | 0 | **+ 52** |
| $\geq 1$ | 1 | **+ 55** |
| $\geq 1$ | 2 | **+ 39** |
| high-confidence HDL particle proteins |  | **296** |
| HDL core proteome: identified in $\geq 80\%$ of all library samples | | **37** |

Comparing our HDL proteome spectral library with the aggregated findings from the other studies, we detected 150 HDL proteins previously identified with high confidence (identified in three or more previous studies). Additionally, we could confirm 94 proteins that were previously identified in one or two previous studies (**Table S2**). We did not, however, detect 73 proteins that were previously found to be associated with HDL in three or more publications. This set of proteins includes mainly subpopulations of immunoglobulins and coagulation and complement factor subunits. These proteins may have been lost during HDL isolation using ultracentrifugation. Further, many of the proteins that were not detected in our analysis contain highly redundant amino acid sequences, so proteotypic peptide-to-protein annotation might have led to association of the peptides we identified with different family members or subunits.

We identified 52 proteins at least twice that had not been associated with HDL in any of the previously published HDL proteomics studies. Although some of these protein identifications might represent isolation artifacts (for instance, proteins co-isolated in microvesicles or exosomes (33) during ultracentrifugation), 13 of these 52 proteins are associated with HDL-mediated functions using GO umbrella terms (**Fig. 2, Table S3**). Among the proteins detected in our analysis, 37 were detected in at least 80% of the samples analyzed. This group is considered to constitute the HDL core proteome. Each of these proteins are also part of the literature-derived HDL proteome. **Table 1** summarizes the numbers of protein groups identified in our study and previous literature.

**Fig. 2.**
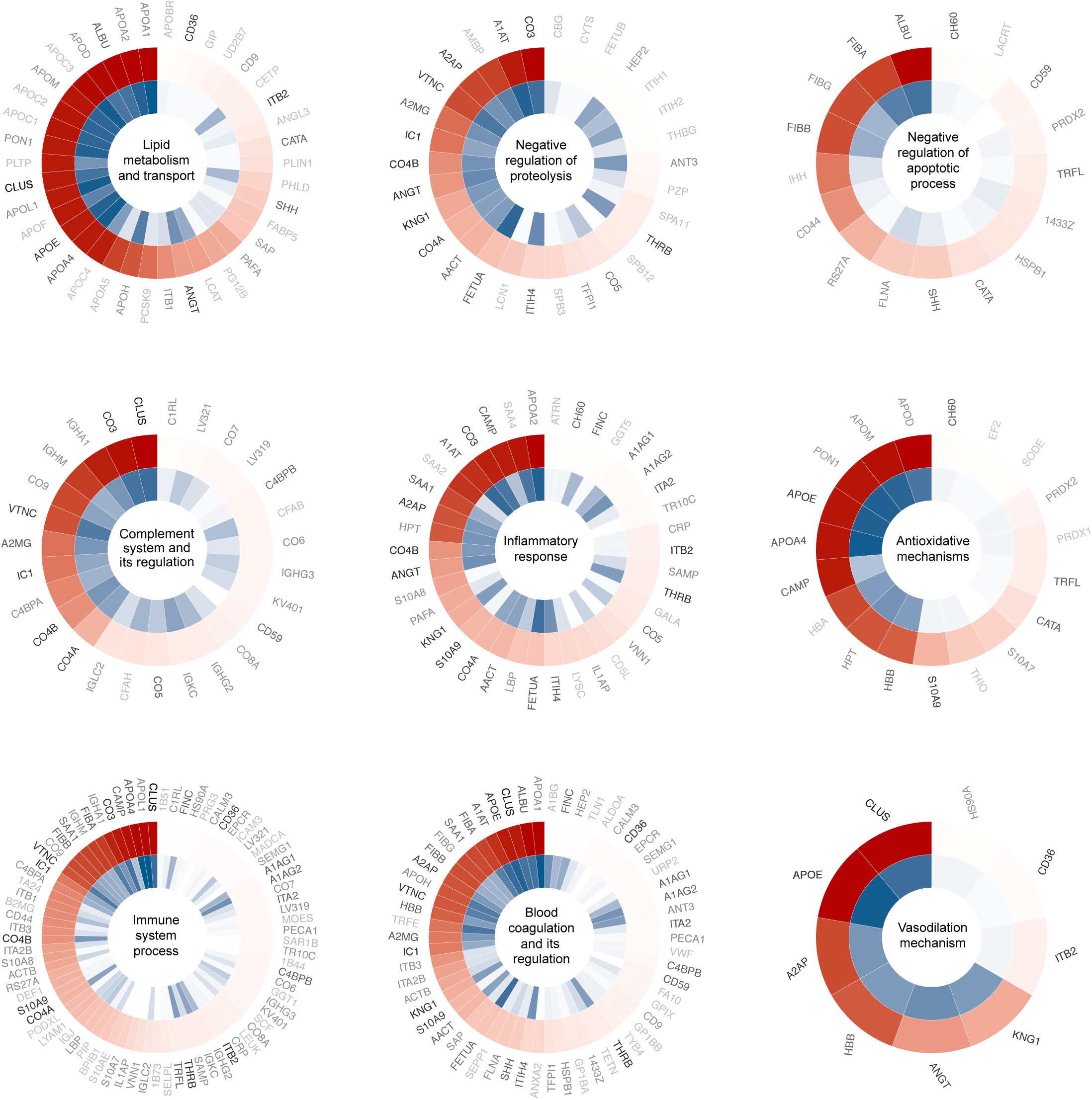
Functional annotation of HDL proteome spectral library proteins. GO analysis grouped the HDL proteome spectral library proteins under nine umbrella terms. HDL proteome spectral library proteins with multiple functional annotations are indicated in black, whereas proteins that are associated with single GO terms are in grey. The outer circle in red shows proteins identified in our analysis (red); the inner circle shows those derived from literature (blue). Color gradients relate to protein group identification frequency (see Fig. 1).

### FUNCTIONAL ANNOTATION OF HDL PROTEOME SPECTRAL LIBRARY PROTEINS

To functionally annotate and classify our high-confidence HDL proteome spectral library proteins, we mapped them to Gene Ontology (GO) biological processes (**Fig. 2**). The GO terms enriched within the proteins from our HDL proteome spectral library fall within nine umbrella terms all associated with known HDL-associated functions. Of the 296 proteins in the spectral library, 171 (58%) were linked to at least one of these umbrella terms. 77 proteins were associated with immune system processes. The rest were associated with blood coagulation and its regulation (57 proteins), lipid metabolism and transport (37 proteins), inflammatory response (38 proteins), negative regulation of proteolysis (30 proteins), complement system and its regulation (27 proteins), antioxidative mechanisms (19 proteins), negative regulation of apoptotic process (16 proteins), and vasodilating mechanisms (9 proteins) (**Table S3**). More than half of the HDL proteome spectral library proteins, including clusterin (CLUS) and apolipoprotein E (APOE) had multifunctional annotations. These proteins might have context-dependent functions related to HDL intrinsic or extrinsic interaction partners. Thus, the results of the functional analysis of the proteins in the HDL proteome spectral library are indicative of a wide range of signaling capacities that could mediate long range signaling functions of HDL particles.

### HDL PROTEOTYPE DIFFERENCES IN BETWEEN HEALTHY SUBJECTS AND THOSE WITH T2DM, CHD, OR T2DM AND CHD

We next quantitatively interrogated HDL proteotype differences in between healthy individuals and subjects with T2DM, CHD, or both T2DM and CHD using the DIA-MS-based HDL proteotyping strategy. Of the 296 proteins in the spectral library, 185 could be quantified across the clinical HDL samples. Statistical analysis revealed significant abundance differences in HDL-associated protein constituents in the patient groups versus the healthy control population (**Fig. 3A, Table S4**). HDL-associated proteins that differed significantly in abundance between the patient groups and the healthy donors were evenly distributed over the whole HDL protein abundance range. This suggests that low abundance HDL proteins, as well as those of high abundance, might be of physiological relevance. (**Fig. 3B, Table S5**). More proteins were downregulated in HDL of patients with T2DM and both T2DM and CHD than were downregulated in patients with CHD. HDL from both T2DM and CHD patients had lower levels of paraoxonases PON1 and PON3 than did HDL of healthy subjects. HDL-associated PONs have atheroprotective effects, by inactivating oxidized lipids, especially lactones (2). Inflammatory proteins, such as serum amyloid A proteins (SAAs), were found present at higher abundance in HDL from subjects with CHD and with both CHD and T2DM than in HDL from healthy subjects, and SAA2 was found to be enriched in all three disease conditions. SAAs interfere with cholesterol efflux capacity (20). Enrichment of HDL with SAA was previously described as an adverse prognostic marker in CHD patients (34). HDL samples from all three patient groups also showed an enrichment of the pulmonary surfactant protein B (PSPB). Longitudinal studies have shown that PSPB-enriched HDL is a significant adverse predictor of mortality in T2DM patients undergoing hemodialysis as well as in patients with heart failure (7,35). Eight other proteins were increased in abundance in patients with disease compared to controls, and six proteins were decreased in abundance across all three disease conditions including PHLD (**Fig. 3C, D and Table S4**).

**Fig. 3.**
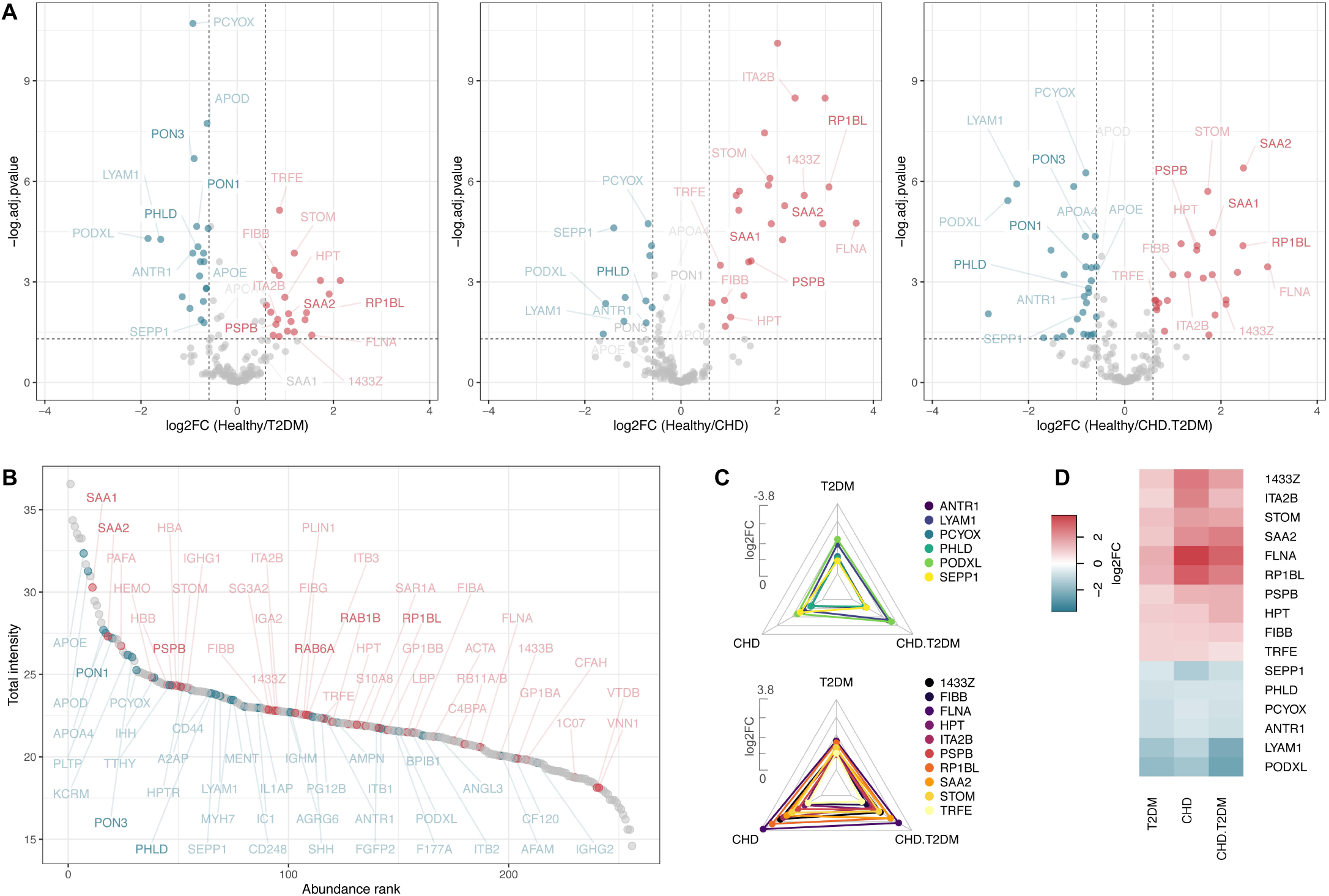
Differences in the HDL proteotypes of healthy individuals and subjects with T2DM, CHD, and both T2DM and CHD. (A) Quantitative HDL proteotype comparison between disease conditions (T2DM, CHD, CHD+T2DM) and a healthy donor cohort. (B) Protein abundance ranking plot of detected HDL proteins over all DIA-quantified samples. Proteins significantly downregulated in pathological conditions are indicated in blue and those significantly upregulated are indicated in red. (C) Radar plots of fold changes derived from the quantitative comparison of HDL from subjects with disease versus healthy controls. Only proteins significantly different in all three conditions are included to show trends over all three comparisons. Upper panel: Proteins depleted in pathological HDL. Lower panel: Proteins enriched in pathological HDL. (D) Protein abundance differences visualized in a heatmap.

### POST-TRANSLATIONAL MODIFICATIONS DETECTED IN HDL PROTEINS AND PATIENT STRATIFICATION BASED ON PEPTIDOFORMS

Similar to other proteins of the human proteome, HDL proteins can be post-translationally modified. These modifications affect structure-function relationships, signaling capacity, and – from a biomedical perspective – the clinical phenotype of patients. Therefore, we analyzed the generated HDL proteotype data using recently developed state-of-the-art bioinformatic tools that enable the detection of peptidoforms without prior enrichment across sample cohorts. Based on an open modification peptide search algorithm (30), we detected 2134 peptides with a q-value *≤* 0.01 (Table S6). 839 out of 2134 peptides were found to be modified including chemical deamidation or carbamidomethylation. 341 peptides of 64 HDL proteins were oxidized (M,W), phosphorylated (S,T,Y), methylated (K), dimethylated (K), or N-terminally acetylated. The most prevalent post-translational modification detected was oxidation of methionine followed by oxidation of tryptophan and phosphorylation of threonine (**Fig. 4A**). The protein with the highest total modified peptidoform signal intensity was APOA1, followed by APOC3 and SAA4 (**Figs. 4B-D, Fig. S1**). APOA1 was previously found to be oxidatively modified at W96 as well as W74 and W132 by myeloperoxidase (36). These modifications were detected in our peptidoform library together with two novel tryptophan oxidations at W22 and W32. Two previously described oxidized methionine residues, M110 and M136, were detected (36) as well as oxidation of M172.

**Fig. 4.**
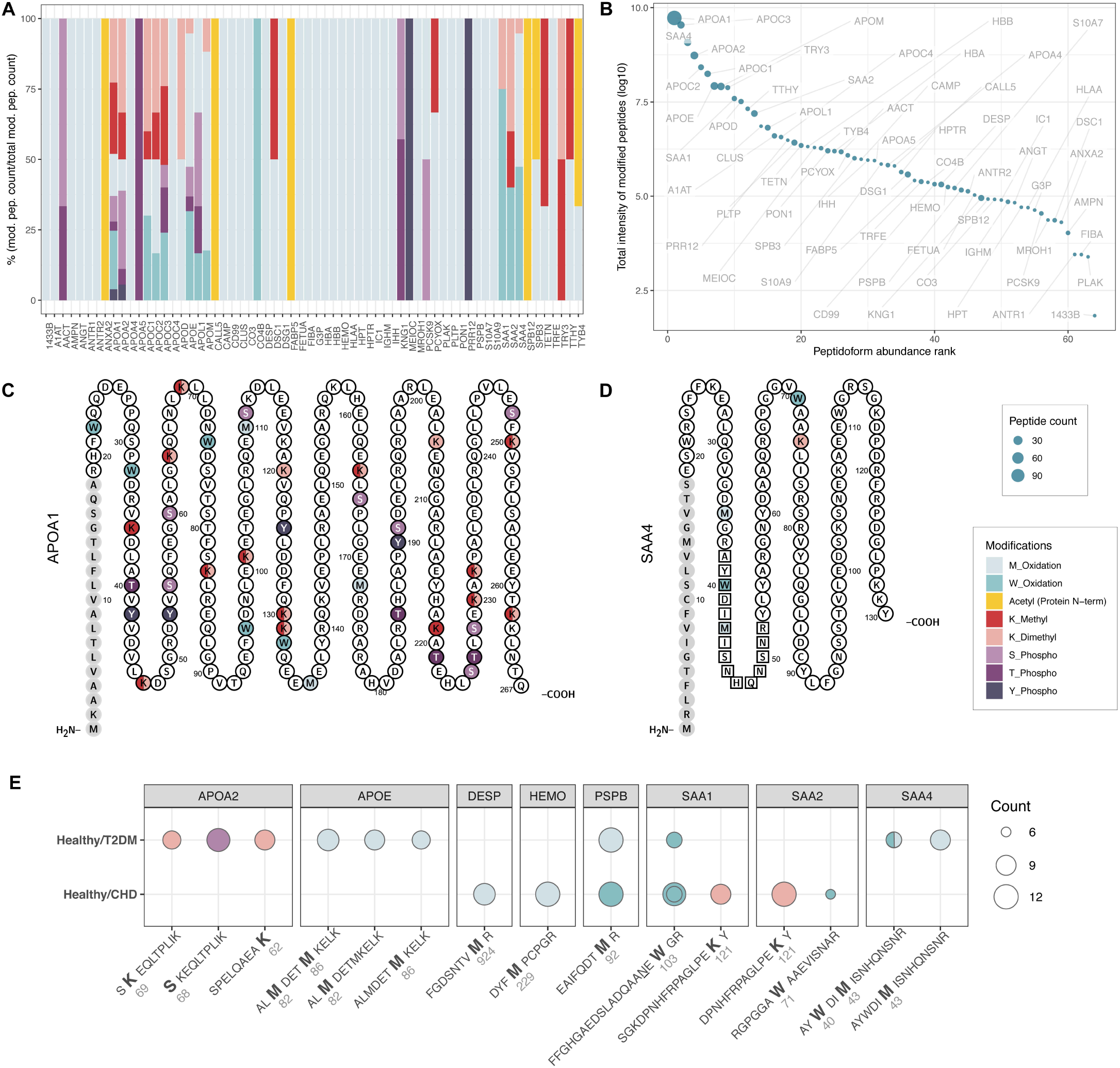
HDL particle peptidoform analysis. (A) Percentages of modified peptidoforms per protein for patient cohorts. Peptides with multiple modifications were treated as separate features. (B) Modified protein intensities derived from the DIA analyses were summed, normalized to peptide length, and plotted according to abundance. The dot size indicates the number of modified peptidoforms per protein. (C,D) Identified amino acid modifications in APOA1 and SAA4. Methyl and dimethyl modifications observed on the same K are shown as semicircles. The (cleaved) signal peptide is shown in grey. The SAA4 peptide depicted in panel E is indicated by squares (http://wlab.ethz.ch/protter). (E) Healthy versus CHD and healthy versus T2DM stratification based on modified peptide intensities normalized to total peptide abundance. Circle size corresponds to the number of iterations in which the feature was selected in the RFE approach. The double circle for oxidized FFGHGAEDSLADQAA<u>N</u>E**W**GR (healthy versus CHD) represents the chemically deamidated peptidoform of the peptide (selected 11 times) and the peptidoform of the same peptide without chemical deamidation (selected 7 times).

Next, we applied the peptidoform information to stratify data from patients based on modified peptide intensities normalized for total peptide abundance. For variable ranking and selection we applied a stratified K-Fold cross (6-fold) validated recursive feature elimination (RFE) approach repeated twice. This process is based on an extremely randomized tree estimator: One thousand trees were used and the RFEs were run until only 10 features remained (**Fig. S2, Table S7**). Several peptidoforms were repetitively selected across the cross-validation runs (**Fig. 4E**). For example, an oxidation motif on methionine at M92 in PSPB, which was previously not characterized, discriminated HDL of both CHD and T2DM patients from healthy subjects. Another interesting example is the downregulation of phosphorylation of a serine residue (S68) in APOA2 in HDL of patients with T2DM. This serine was previously identified as the target of the Fam20C kinase, which is involved in lipid homeostasis (37). Overall, we observed that quantitative peptide information, which is related to protein quantity, best stratifies different patient groups, whereas peptide modifications are minor discriminators between pathological HDL phenotypes (**Fig. S2**).

## Discussion

HDL is a complex blood-circulating mix of particles varying by size and molecular composition. Despite its acknowledged importance for many biological functions, structure-function relationships are largely not resolved yet – especially with regard to various pathologies. To link structure with function one needs to acquire consistent information about the constituents of HDL particles. Here we developed a strategy and toolbox which enables sensitive and reproducible HDL analysis on the proteotype level, which is fundamental for clinical analysis.

Our DIA-MS analysis approach not only enables the digitization of HDL patient cohorts, but also a multi-centric, distributed data generation and analysis as eventually required for observational and interventional clinical trials (29).

The HDL proteome spectral libraries we developed can serve as a rather advanced starting point and community reference for the analytical interrogation of HDL in healthy individuals and patients. Spectral information is provided for more than 4000 peptides and peptidoforms and enables the sensitive, reproducible and relative quantitative analysis of 296 high-confident HDL proteins. Peptide spectral library matching, as opposed to standard sequence matching to fragmentation spectra generated *in silico*, typically provides greater sensitivity and specificity for peptide identification. This is of particular importance for DIA, where highly complex and multiplexed fragmentation spectra are acquired. HDL peptidoforms carrying biologically relevant post-translational modifications are often more difficult to measure consistently across larger sample cohorts as they are frequently less stable or less abundant than unmodified peptides. Due to the acquisition of full fragment ion spectra, DIA also provides an ideal platform for the efficient and accurate quantitation of peptidoforms and pathological PTMs across large HDL sample cohorts. Here we demonstrated that a peptidoform-based DIA HDL quantitation strategy enabled the identification of novel determinants of pathological HDL, and future applications of this strategy to larger HDL cohorts could help to unravel HDL-structure-function-disease relationships.

We utilized our newly developed HDL proteome spectral library for the relative quantitative analysis of the composition of HDL proteoforms in patients suffering from T2DM and/or CHD. Our data does not immediately enable us to draw conclusions on whether detected molecular alterations are causes, effects, or bystanders of disease conditions; however, we can make assumptions and generate hypotheses on their potential contributions to molecular functions in the human body, such as long range signaling through HDL. The composition of individual HDL particles, which we did not investigate here due to current technological limitations will likely influence molecular interaction capabilities. Based on proteins identified in this and other studies it is conceivable that many different HDL particles of varying constituents exist, which stochastically carry proteoforms. Depending on their co-existence on a single HDL particle a particular function, destination and interaction space could be conferred. The decoding of such a potential HDL barcode is the next analytical challenge in elucidating HDL structure-function relationship.

An interesting finding of our quantitative analysis of the HDL cohorts was the significant depletion of phospholipase PHLD from HDL of all patient groups. PHLD hydrolyzes the glycan-phosphatidylinositol linkages of glycosylphosphatidylinositol-anchored cell surface proteins. We previously showed that PHLD inhibits apoptosis in human aortic endothelial cells, and thereby exerts a presumably vasoprotective function (3). Also, we identified several relatively low-abundant Ras-related proteins that are enriched in HDL particles from patients including RAB6A, RB11A/B, RAB1B, and RP1BL (**Fig. 3B**). Rab proteins are small GTPases that are key regulators of intracellular membrane trafficking. These proteins are components of extracellular vesicles (33). Therefore and because of overlapping density, we cannot exclude that our HDL isolates were contaminated with exosomes or microvesicular bodies which contain among other proteins also Rab proteins. However, it may also be that HDL associates with Rab proteins during transcytosis or retroendocytosis through various cell types including endothelial cells, macrophages or hepatocytes (38). Tubular endosome formation in return is regulated by a number of Rab family members (39). It is therefore conceivable that some of these RAB proteins localize to HDL during intracellular trafficking. Also, Rab-dependent membrane transport functions in clearance of cholesterol from late endosomes (40). Recently, RAB10 was identified in HDL from patients suffering from chronic heart failure (15).

In summary, the DIA-based HDL proteotyping strategy developed here enables now sensitive and reproducible digitization of HDL proteotypes and provides new insights into the composition of HDL particles as a rational basis for decoding clinically relevant HDL structure-function relationships.

## Acknowledgments

We thank J. R. Wyatt for text editing and the staff of the Clinical Trial Center at USZ for the recruitment and management of healthy volunteers and patients. We thank Dr. Wenguang Shao for bioinformatic expertise in analyzing data with MsFragger.

This work was supported by ETH grant 30 17-1 (to B.W.), the Swiss National Science Foundation (grant 31003A_160259 to B.W), a SystemsX.ch special opportunity grant (to B.W.), a SystemsX.ch MRD grant (2014/267 to A.v.E. and B.W.) and the FP7 Project “RESOLVE” (to A.v.E.).

## Author contributions

SG and KF performed all experiments and analyses except those noted below. LR and SR managed the biobank and isolated HDL. AVE, JK, UL, and MB obtained ethical approval for the studies and recruited patients and healthy volunteers. PGAP performed quantitative peptidoform analysis and provided bioinformatic expertise. SG, KF, and BW designed research. AVE and LR provided expertise and critical feedback at all stages of the project. SG, KF, and BW conceived the project and wrote the paper. All authors critically read and revised the manuscript.

## List of abbreviations

HDL: high-density lipoprotein
NO: nitric oxide
HDL-C: high-density lipoprotein cholesterol
CHD: coronary heart disease
T2DM: diabetes mellitus type 2
DDA: data-dependent acquisition
DIA: data-independent acquisition
MS: mass spectrometry
PTMs: post-translational modifications
GO: Gene Ontology
RFE: recursive feature elimination

## References

1. Rosenson RS, Brewer HB Jr, Davidson WS, Fayad ZA, Fuster V, Goldstein J, et al. Cholesterol efflux and atheroprotection: advancing the concept of reverse cholesterol transport. Circulation. 2012;125:1905–19.

2. Besler C, Lüscher TF, Landmesser U. Molecular mechanisms of vascular effects of High-density lipoprotein: alterations in cardiovascular disease. EMBO Mol Med. 2012;4:251–68.

3. Cardner M, Yalcinkaya M, Goetze S, Luca E, Balaz M, Hunjadi M, et al. Structure-function relationships of HDL in diabetes and coronary heart disease. JCI Insight [Internet]. 2019; Available from: http://dx.doi.org/10.1172/jci.insight.131491

4. März W, Kleber ME, Scharnagl H, Speer T, Zewinger S, Ritsch A, et al. HDL cholesterol: reappraisal of its clinical relevance [Internet]. Clinical Research in Cardiology. 2017. page 663–75. Available from: http://dx.doi.org/10.1007/s00392-017-1106-1

5. Burillo E, Jorge I, Martínez-López D, Camafeita E, Blanco-Colio LM, Trevisan-Herraz M, et al. Quantitative HDL Proteomics Identifies Peroxiredoxin-6 as a Biomarker of Human Abdominal Aortic Aneurysm. Sci Rep. 2016;6:38477.

6. Collier TS, Jin Z, Topbas C, Bystrom C. Rapid Affinity Enrichment of Human Apolipoprotein A-I Associated Lipoproteins for Proteome Analysis. J Proteome Res. 2018;17:1183–93.

7. Emmens JE, Jones DJL, Cao TH, Chan DCS, Romaine SPR, Quinn PA, et al. Proteomic diversity of high-density lipoprotein explains its association with clinical outcome in patients with heart failure. Eur J Heart Fail. 2018;20:260–7.

8. Fournier M, Bonneil E, Garofalo C, Grimard G, Laverdière C, Krajinovic M, et al. Altered proteome of high-density lipoproteins from paediatric acute lymphoblastic leukemia survivors. Sci Rep. 2019;9:4268.

9. Godzien J, Ciborowski M, Armitage EG, Jorge I, Camafeita E, Burillo E, et al. A Single In-Vial Dual Extraction Strategy for the Simultaneous Lipidomics and Proteomics Analysis of HDL and LDL Fractions. J Proteome Res. 2016;15:1762–75.

10. Gordon SM, McKenzie B, Kemeh G, Sampson M, Perl S, Young NS, et al. Rosuvastatin Alters the Proteome of High Density Lipoproteins: Generation of alpha-1-antitrypsin Enriched Particles with Anti-inflammatory Properties. Mol Cell Proteomics. 2015;14:3247–57.

11. Gourgari E, Ma J, Playford MP, Mehta NN, Goldman R, Remaley AT, et al. Proteomic alterations of HDL in youth with type 1 diabetes and their associations with glycemic control: a case-control study. Cardiovasc Diabetol. 2019;18:43.

12. Ljunggren SA, Helmfrid I, Norinder U, Fredriksson M, Wingren G, Karlsson H, et al. Alterations in high-density lipoprotein proteome and function associated with persistent organic pollutants. Environ Int. 2017;98:204–11.

13. Mathew AV, Li L, Byun J, Guo Y, Michailidis G, Jaiswal M, et al. Therapeutic Lifestyle Changes Improve HDL Function by Inhibiting Myeloperoxidase-Mediated Oxidation in Patients With Metabolic Syndrome. Diabetes Care. 2018;41:2431–7.

14. Melchior JT, Street SE, Andraski AB, Furtado JD, Sacks FM, Shute RL, et al. Apolipoprotein A-II alters the proteome of human lipoproteins and enhances cholesterol efflux from ABCA1. J Lipid Res. 2017;58:1374–85.

15. Oberbach A, Adams V, Schlichting N, Heinrich M, Kullnick Y, Lehmann S, et al. Proteome profiles of HDL particles of patients with chronic heart failure are associated with immune response and also include bacteria proteins. Clin Chim Acta. 2016;453:114–22.

16. Okada T, Ohama T, Takafuji K, Kanno K, Matsuda H, Sairyo M, et al. Shotgun proteomic analysis reveals proteome alterations in HDL of patients with cholesteryl ester transfer protein deficiency. J Clin Lipidol. 2019;13:317–25.

17. Pedret A, Catalán Ú, Fernández-Castillejo S, Farràs M, Valls R-M, Rubió L, et al. Impact of Virgin Olive Oil and Phenol-Enriched Virgin Olive Oils on the HDL Proteome in Hypercholesterolemic Subjects: A Double Blind, Randomized, Controlled, Cross-Over Clinical Trial (VOHF Study). PLoS One. 2015;10:e0129160.

18. Rao PK, Merath K, Drigalenko E, Jadhav AYL, Komorowski RA, Goldblatt MI, et al. Proteomic characterization of high-density lipoprotein particles in patients with non-alcoholic fatty liver disease. Clin Proteomics. 2018;15:10.

19. Swertfeger DK, Li H, Rebholz S, Zhu X, Shah AS, Davidson WS, et al. Mapping Atheroprotective Functions and Related Proteins/Lipoproteins in Size Fractionated Human Plasma. Mol Cell Proteomics. 2017;16:680–93.

20. Vaisar T, Tang C, Babenko I, Hutchins P, Wimberger J, Suffredini AF, et al. Inflammatory remodeling of the HDL proteome impairs cholesterol efflux capacity. J Lipid Res. 2015;56:1519–30.

21. Gordon SM, Chung JH, Playford MP, Dey AK, Sviridov D, Seifuddin F, et al. High density lipoprotein proteome is associated with cardiovascular risk factors and atherosclerosis burden as evaluated by coronary CT angiography. Atherosclerosis. 2018;278:278–85.

22. Silva ARM, Toyoshima MTK, Passarelli M, Di Mascio P, Ronsein GE. Comparing Data-Independent Acquisition and Parallel Reaction Monitoring in Their Abilities To Differentiate High-Density Lipoprotein Subclasses. J Proteome Res [Internet]. 2019; Available from: http://dx.doi.org/10.1021/acs.jproteome.9b00511

23. Florens N, Calzada C, Delolme F, Page A, Guebre Egziabher F, Juillard L, et al. Proteomic Characterization of High-Density Lipoprotein Particles from Non-Diabetic Hemodialysis Patients. Toxins [Internet]. 2019;11. Available from: http://dx.doi.org/10.3390/toxins11110671

24. Ronsein GE, Vaisar T. Deepening our understanding of HDL proteome. Expert Rev Proteomics [Internet]. 2019; Available from: http://dx.doi.org/10.1080/14789450.2019.1650645

25. Charles-Schoeman C, Gugiu GB, Ge H, Shahbazian A, Lee YY, Wang X, et al. Remodeling of the HDL proteome with treatment response to abatacept or adalimumab in the AMPLE trial of patients with rheumatoid arthritis. Atherosclerosis. 2018;275:107–14.

26. Sean Davidson W, Silva RAGD, Chantepie S, Lagor WR, John Chapman M, Kontush A. Proteomic Analysis of Defined HDL Subpopulations Reveals Particle-Specific Protein Clusters. Arterioscler Thromb Vasc Biol. American Heart Association, Inc.; 2009;29:870–6.

27. Heinecke JW. The HDL proteome: a marker–and perhaps mediator–of coronary artery disease: Fig. 1 [Internet]. Journal of Lipid Research. 2009. page S167–71. Available from: http://dx.doi.org/10.1194/jlr.r800097-jlr200

28. Bruderer R, Bernhardt OM, Gandhi T, Xuan Y, Sondermann J, Schmidt M, et al. Optimization of Experimental Parameters in Data-Independent Mass Spectrometry Significantly Increases Depth and Reproducibility of Results. Mol Cell Proteomics. 2017;16:2296–309.

29. Xuan Y, Bateman NW, Gallien S, Goetze S, Zhou Y, Navarro P, et al. Standardization and Harmonization of Distributed Multi-National Proteotype Analysis supporting Precision Medicine Studies [Internet]. Available from: http://dx.doi.org/10.1101/2020.03.12.988089

30. Kong AT, Leprevost FV, Avtonomov DM, Mellacheruvu D, Nesvizhskii AI. MSFragger: ultrafast and comprehensive peptide identification in mass spectrometry-based proteomics. Nat Methods. 2017;14:513–20.

31. Choi M, Chang C-Y, Clough T, Broudy D, Killeen T, MacLean B, et al. MSstats: an R package for statistical analysis of quantitative mass spectrometry-based proteomic experiments. Bioinformatics. 2014;30:2524–6.

32. Pedregosa F, Varoquaux G, Gramfort A, Michel V, Thirion B, Grisel O, et al. Scikit-learn: Machine Learning in Python. J Mach Learn Res. 2011;12:2825–30.

33. Karimi N, Cvjetkovic A, Jang SC, Crescitelli R, Hosseinpour Feizi MA, Nieuwland R, et al. Detailed analysis of the plasma extracellular vesicle proteome after separation from lipoproteins. Cell Mol Life Sci. 2018;75:2873–86.

34. Zewinger S, Drechsler C, Kleber ME, Dressel A, Riffel J, Triem S, et al. Serum amyloid A: high-density lipoproteins interaction and cardiovascular risk. Eur Heart J. 2015;36:3007–16.

35. Kopecky C, Genser B, Drechsler C, Krane V, Kaltenecker CC, Hengstschläger M, et al. Quantification of HDL proteins, cardiac events, and mortality in patients with type 2 diabetes on hemodialysis. Clin J Am Soc Nephrol. 2015;10:224–31.

36. Rosenberger G, Liu Y, Röst HL, Ludwig C, Buil A, Bensimon A, et al. Inference and quantification of peptidoforms in large sample cohorts by SWATH-MS. Nat Biotechnol [Internet]. 2017; Available from: http://dx.doi.org/10.1038/nbt.3908

37. Tagliabracci VS, Wiley SE, Guo X, Kinch LN, Durrant E, Wen J, et al. A Single Kinase Generates the Majority of the Secreted Phosphoproteome. Cell. 2015;161:1619–32.

38. Zanoni P, Velagapudi S, Yalcinkaya M, Rohrer L, von Eckardstein A. Endocytosis of lipoproteins. Atherosclerosis. 2018;275:273–95.

39. Etoh K, Fukuda M. Rab10 regulates tubular endosome formation through KIF13A and KIF13B motors. J Cell Sci [Internet]. 2019;132. Available from: http://dx.doi.org/10.1242/jcs.226977

40. Ahras M, Naing T, McPherson R. Scavenger receptor class B type I localizes to a late endosomal compartment. J Lipid Res. 2008;49:1569–76.

