## Supplemental Data for "Sensitive and reproducible determination of clinical HDL proteotypes"

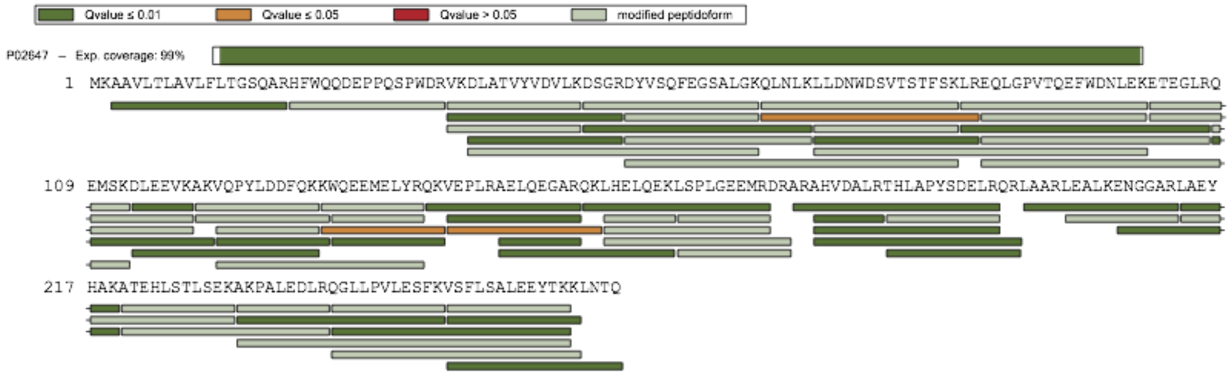


**Figure S1: Sequence coverage of APOPA1.**

APOA1 was identified with 99% sequence coverage. Stripped peptide sequences identified with high confidence are displayed as green bars. Modified peptide sequences are shown in light green.

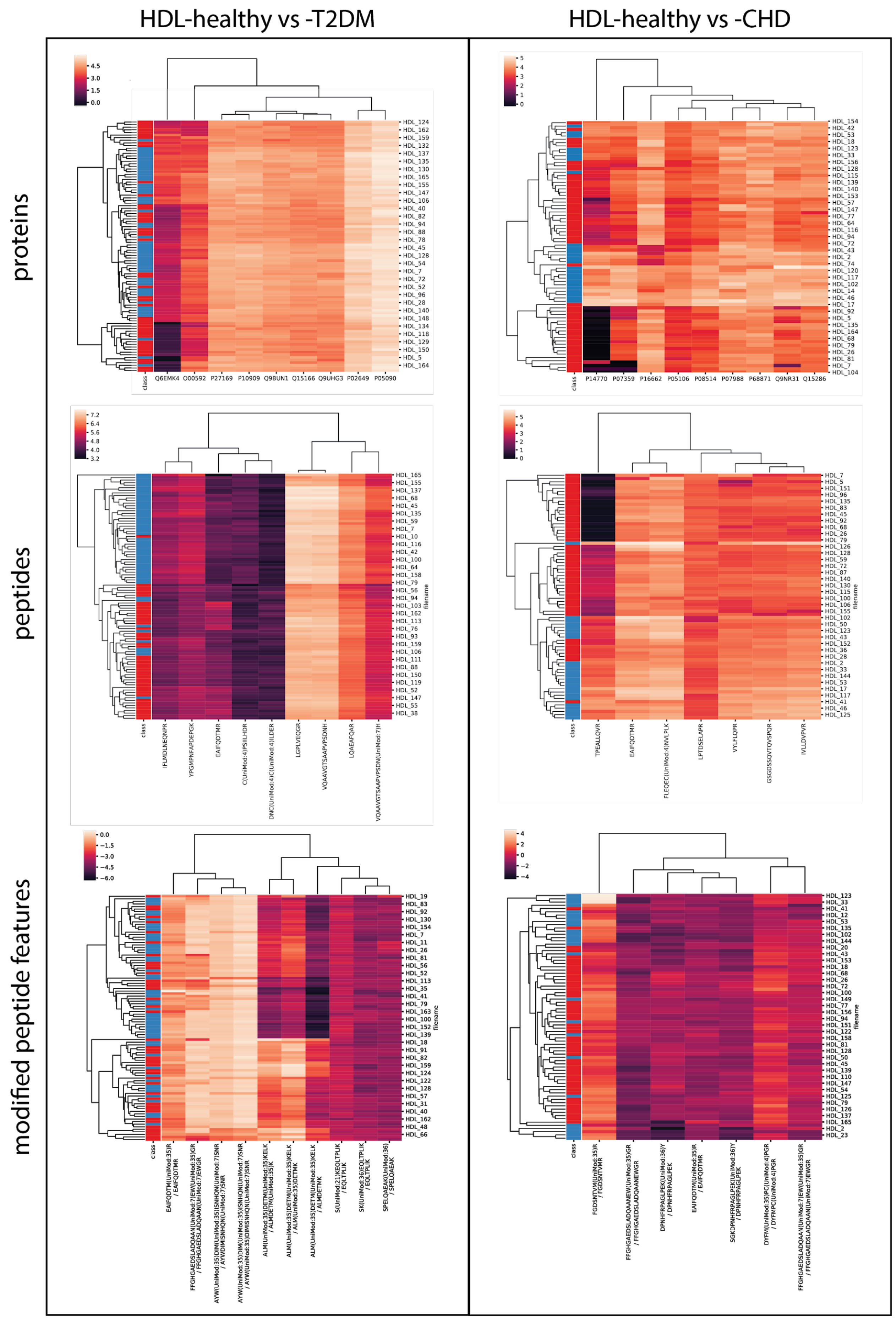

HDL-healthy vs -T2DM
HDL-healthy vs -CHD
proteins
peptides
modified peptide features
class
Q6EMK4 O00592 P27169 P10909 Q98UN1 Q15166 Q9UHG3 P02649 P05090
P14770 P07359 P16662 P05106 P08514 P07988 P68871 Q9NR31 Q15286
filename

**Figure S2: Clustering by recursively selected features.**

The 10 most important features were repeatedly selected (scikitlearn RepeatedStratifiedKFold, 2 repeats and 6 folds) using recursive feature elimination using an ExtraTreesClassifier scikitlearn estimator. Features selected in more than half of the cross-validation runs were used to generate the clustermaps. Clustering by protein, peptide, and modified peptide feature ratios are shown for HDL from healthy subjects versus HDL from T2DM (left) and CHD (right) patients. Red: diseased; blue: healthy

**Table S1** - Summary of all identified protein groups (including HDL-related contaminants).
**Table S2** - HDL particle proteins identified in two or more measurements, in one measurement with additional evidence from literature, or with literature evidence from three or more studies.
**Table S3** - Gene Ontology biological processes umbrella term associations of HDL particle proteins.
**Table S4** - Quantitative comparisons of components of HDL particles from patients with CHD, T2DM, or both diseases relative to components from healthy subjects (results of MSstats significance testing).
**Table S5** - DIA protein quantification of HDL isolates (unfiltered).
**Table S6 -** DIA PTM localization probabilities and PTM quantification of HDL isolates (unfiltered).
**Table S7 -** Top discriminating PTM features based on *k*-nearest-neighbours analysis.
**Table S8** - Overview of DIA scan windows.
