## Supplemental Materials and Methods for "Sensitive and reproducible determination of clinical HDL proteotypes"

### **Clinical chemistry measurements of plasma samples**

Activities of liver enzymes aspartate transaminase and alanine transaminase as well as concentrations of total and HDL cholesterol, triglycerides, glucose, creatinine, glycated hemoglobin, and c-reactive protein were measured using a COBAS8000 analyser and assays from Roche diagnostics (Rotkreuz Switzerland) in the of the Institute of Clinical Chemistry, University Hospital of Zurich.

### **HDL sample preparation**

HDL particles were isolated by differential ultracentrifugation basically as described by Havel et al. (1). We adapted the original protocol depending on the sample volume and applied the two different protocols to isolate HDL. In all cases, the density was adjusted to 1.063 with KBr, samples were centrifuged, and the top layer containing LDL particles was removed. Subsequently, the density was raised to 1.21 with KBr, the samples were centrifuged, and the top layer containing HDL was collected. For large volumes (10 mL plasma and more), 1.5 mL saturated KBr solution was put into Quickseal tubes (Beckman 342413), filled to the top with KBr (to d=1.063), and samples were centrifuged at 250,000 x g at 15 °C for 16 h. The tubes were cut 3 cm from the bottom, and upper fractions containing chylomicrons, VLDL, and LDL were discarded. The lower fractions with the HDL were mixed with 5 mL saturated KBr solution, transferred to a new Quickseal tube (Beckman 342413), filled with KBr (to d=1.21) and centrifuged again at 250,000 x g at 15 °C for 16 h. Subsequently, the 3 mL at the top of the tube containing HDL were collected. To the 3-mL HDL-containing fraction was added 1.5 mL saturated KBr solution. The samples was transferred to a new Quickseal tube (Beckman 342413), filled with KBr solution (to d= 1.21), and spun at 250,000 x g at 15 °C for 16 h. The top 600 μL were collected and dialyzed twice against 2 L 150 mM NaCl containing 0.3 mM EDTA , pH 7.4 at 4°C to remove KBr.

For small volumes (0.5 mL), plasma was mixed with 1 mL 150 mM NaCl containing 1 mM EDTA and 0.5 mL saturated KBr (to d=1.32) in a Quickseal tube (Beckman 344625). Samples were centrifuged at 550,000 x g at 4 °C for 6 h. The HDL-containing fraction was collected by cutting the tube approximately 10 mm from the bottom and discarding the upper fraction. The lower fraction with the HDL was adjusted to a density of 1.21 by mixing with 0.9 mL saturated KBr and was centrifuged at 550,000 x g at 4 °C for 16 h. The top fraction containing HDL (about 150 μL) was collected by cutting the tube 15 mm from the bottom. To remove KBr, the collected HDL fractions were concentrated and dialyzed three times against 150 mM NaCl containing 0.3 mM EDTA, pH 7.4 using Amicon Ultra Centrifugal Filters, Ultracel 30K.

### **Sample preparation for mass spectrometry**

HDL was delipidated and processed as described elsewhere (2). In short, 80-100 μg of HDL protein were extracted with methanol/chloroform (2:1). For phase separation 0.3 ml of water were added. The upper phase was discarded and proteins from the interphase were precipitated by the addition of methanol. To monitor extraction efficiency and reproducibility, bovine alpha-1-acid-Glycoprotein (Sigma-Aldrich) was spiked into the samples as an external reference (2 mg/ml final concentration). For one sample set, methyl-tert butyl ether extraction was applied (3). After protein precipitation, pellets were resuspended in 100 𝜇l sodium deoxycholate lysis buffer (1% w/v) in 100 mM Tris-HCl (pH 8.5) containing 10 mM Tris(2-carboxyethyl)phosphine and 15 mM chloroacetamide (all Sigma-Aldrich). Samples were denatured, reduced, and alkylated at 95 °C for 5 min. Samples were treated with sequencing-grade trypsin (Promega) at a ratio of 1:40 for 18 h at 37 °C. Sample purification was performed over C18 resin (100 𝜇g capacity columns or plates; The Nest Group) using 2% to 80% acetonitrile, 0.1% formic acid (FA; Chemie Brunschwig). For mass spectrometry analysis, we injected 2 μg peptides per sample together with retention time reference peptides (iRTs; Biogosys) on a 50-cm column.

### **DDA mass spectrometry measurements for spectral library generation**

Peptide samples were separated by reversed-phase chromatography (EASY-Spray RSLC C18, 2 𝜇m, 50 cm x 75 𝜇m; Thermo Fisher Scientific), which was connected to a nano-flow HPLC interfaced with an autosampler (EASY-nLC 1200, Proxeon). The HPLC was coupled to a QExactive HF (Thermo Fisher Scientific) equipped with an Easy-Spray ion source (Thermo Fisher Scientific). Peptides were loaded onto a trap column (C18, 3 𝜇m, 75 𝜇m x 20 mm; Thermo Fisher Scientific) with 100% buffer A (99.9% H_2_O, 0.1% FA) and eluted onto a 50-cm Easy-Spray PepMap C18 column (ES803, 2 𝜇m, 75 𝜇m x 50 cm) (Thermo Fisher Scientific) at 50 °C with a constant flow rate of 200 nl/min with a 30-min linear gradient from 5–32% buffer B (80% acetonitrile, 0.1% FA) and 5 min 32-56% buffer B. After the gradient, the column was washed twice with 100% buffer B. Mass spectra were acquired in a DDA measurement mode. High-resolution MS1 spectra were acquired at 60,000 resolution (automatic gain control target value 3 x 10^6^) to monitor peptide ions in the mass range of 375–1,500 m/z, followed by HCD MS/MS scans at 15,000 resolution (automatic gain control target value 1 x 10^5^). Dynamic exclusion was set to 15 s. MS/MS were recorded in centroid mode.

### **DDA data processing details for spectral library generation**

For spectral library generation, RAW files were processed with Proteome Discoverer software, version 2.2 using a human UniProt database (release April 2018) together with iRT peptides and common contaminants. The processing workflow consisted of SequestHT (4) and Amanda (5) search nodes coupled with Percolator (6). The following search parameters were used for protein identification: (i) peptide mass tolerance set to 10 ppm; (ii) MS/MS mass tolerance set to 0.02 Da; (iii) fully tryptic peptides with up to two missed cleavages were allowed; (iv) carbamidomethylation of cysteine was set as fixed modification; methionine oxidation, serine/threonine/tyrosine phosphorylation, asparagine deamidation, and protein N-terminal acetylation were set as variable modifications. For our peptidoform library, we additionally included tryptophan oxidation as well as mono- and di-methylation of lysine as variable modifications. Percolator was set at max delta Cn 0.05, with target FDR strict 0.01 and target FDR relaxed 0.05.

### **DIA Mass Spectrometry Measurements and Analysis of clinical cohort**

For DIA, the same gradient used for DDA was applied. The DIA-MS method consisted of one MS1 scan from 375 to 1200 m/z at 60,000 resolution (AGC target 3 x 10^6^, maximum IT 55ms), followed by 19 consecutive DIA segments acquired at 30,000 resolution (AGC target 3 x 10e^6^, maximum IT auto). DIA segments were recorded in profile mode with variable overlapping scan windows (**Table S8**). Normalized collision energy was stepped between 25 and 30. The spectra were recorded in profile mode.

DIA data were analysed using Spectronaut in default mode. For retention time calibration, iRTs (Biognosys) were added to each sample. For quantitation, standard settings were employed, which included dynamic peak detection, automatic precision nonlinear iRT calibration, interference correction, local cross run normalization, and MS2 Top 3 summed peptide quantitation. All results were filtered with a q-value of 0.01 (equal to an FDR of 1%) on the peptide level. The data matrix was filtered with q-value sparse (i.e., all observations that pass the q-value threshold of 0.01 at least once were considered). Data files were searched against the human UniProt fasta database (release April 2018). Confidence on the peptidoform-level was evaluated using an FDR of 1%. For peptidoform visualization on the protein level, Protter was used (7).

For statistical data evaluation, MSstats3 (v3.12.3) was used (8). Normalization was chosen to be quantile, imputation was selected. The other parameters were set to default. A minimum of four features (peptideSequence, chargeState, fragmentIon, combination) was selected per protein and run. Tests for significant changes in protein abundance across conditions were based on a family of linear mixed-effects models. The p-values were multiple testing corrected to control the experiment-wide FDR at a desired level using the Benjamini-Hochberg method. Proteins were considered differentially expressed if they showed a fold-change of 1.5 or higher and an adjusted p-value of 0.05 or lower.

For modified peptide selection and feature clustering, the quantitative peptide matrix generated by Spectronaut based on the peptidoform library with additional PTMs was processed by removing mass spectrometry contaminants and non-proteotypic peptides. After converting intensities to log10 and summing values for alternative charge states, peptides with log10(intensity) values ≤ 2.5 across all conditions were removed. Furthermore, samples with ≥ 70% and peptides with ≥ 90% missing data values were removed. The filtered quantitative matrix was median normalized and missing values were filled in with the minimum value of the feature across all samples minus a random 1-10%. At this point the data strongly clustered in the first principal component dimension with a technical sample preparation factor (ultra-centrifugation batch). To remove this batch effect the first principal component dimension was dropped as well as two samples deemed to be outliers (HDL 16 and 22).

1.	Havel RJ, Eder HA, Bragdon JH. The distribution and chemical composition of ultracentrifugally separated lipoproteins in human serum. J Clin Invest. 1955;34:1345–53.

2.	[Cardner M, Yalcinkaya M, Goetze S, Luca E, Balaz M, Hunjadi M, et al. Structure-function relationships of HDL in diabetes and coronary heart disease. JCI Insight [Internet]. 2019; Available from:](http://paperpile.com/b/Up5UIA/rny6b) <http://dx.doi.org/10.1172/jci.insight.131491>

3.	Matyash V, Liebisch G, Kurzchalia TV, Shevchenko A, Schwudke D. Lipid extraction by methyl-tert-butyl ether for high-throughput lipidomics. J Lipid Res. 2008;49:1137–46.

4.	Eng JK, McCormack AL, Yates JR. An approach to correlate tandem mass spectral data of peptides with amino acid sequences in a protein database. J Am Soc Mass Spectrom. 1994;5:976–89.

5.	Dorfer V, Pichler P, Stranzl T, Stadlmann J, Taus T, Winkler S, et al. MS Amanda, a universal identification algorithm optimized for high accuracy tandem mass spectra. J Proteome Res. 2014;13:3679–84.

6.	Brosch M, Yu L, Hubbard T, Choudhary J. Accurate and sensitive peptide identification with Mascot Percolator. J Proteome Res. 2009;8:3176–81.

7.	Omasits U, Ahrens CH, Müller S, Wollscheid B. Protter: interactive protein feature visualization and integration with experimental proteomic data. Bioinformatics. 2014;30:884–6.

8.	Choi M, Chang C-Y, Clough T, Broudy D, Killeen T, MacLean B, et al. MSstats: an R package for statistical analysis of quantitative mass spectrometry-based proteomic experiments. Bioinformatics. 2014;30:2524–6.
